# Parallelization with Dual-Trap Single-Column Configuration Maximizes Throughput of Proteomic Analysis

**DOI:** 10.1101/2022.06.02.494601

**Authors:** Simion Kreimer, Ali Haghani, Aleksandra Binek, Alisse Hauspurg, Saeed Seyedmohammad, Alejandro Rivas, Amanda Momenzadeh, Jesse Meyer, Koen Raedschelders, Jennifer E. Van Eyk

## Abstract

Proteomic analysis on the scale that captures population and biological heterogeneity over hundreds to thousands of samples requires rapid mass spectrometry methods which maximize instrument utilization (IU) and proteome coverage while maintaining precise and reproducible quantification. To achieve this, a short liquid chromatography gradient paired to rapid mass spectrometry data acquisition can be used to reproducibly profile a moderate set of analytes. High throughput profiling at a limited depth is becoming an increasingly utilized strategy for tackling large sample sets but the time spent on loading the sample, flushing the column(s), and re-equilibrating the system reduces the ratio of meaningful data acquired to total operation time and IU. The dual-trap single-column configuration presented here maximizes IU in rapid analysis (15 min per sample) of blood and cell lysates by parallelizing trap column cleaning and sample loading and desalting with analysis of the previous sample. We achieved 90% IU in low micro-flow (9.5 µL/min) analysis of blood while reproducibly quantifying 300-400 proteins and over 6,000 precursor ions. The same IU was achieved for cell lysates, in which over 4,000 proteins (3,000 at CV below 20%) and 40,000 precursor ions were quantified at a rate of 15 minutes/sample. Thus, deployment of this dual-trap single column configuration enables high throughput epidemiological blood-based biomarker cohort studies and cell-based perturbation screening.

## Introduction

Automation of sample preparation and the commercialization of remote sampling devices have overcome two bottlenecks in large scale mass spectrometry based proteomic studies. The once cumbersome sample preparation for bottom-up proteomic analysis in which the protein content is extracted and enzymatically digested into peptides [1] can now reproducibly prepare several hundred to thousands of samples per day through automation [2]. Remote sampling devices allow donors to autonomously and reproducibly extract and ship several microliters of blood, thus greatly accelerating collection of specimens from many cohorts across multiple time points. With these developments, the remaining bottleneck is the analysis itself in which the generated peptides are separated by liquid chromatography and quantified by mass spectrometry (LC-MS). While multiplexing through isobaric mass tag labelling significantly improves LC-MS throughput [3-8], limitations of this strategy across large sample sets [9-11] have encouraged the use of rapid, individual, analysis of samples with data independent acquisition (DIA) [12-14]. The challenge is now to balance the maximum number of reproducibly quantified protein species with the speed of analysis.

Epidemiological biomarker investigations are a key application that benefits from remote sampling and requires high throughput. Blood collected through Mitra devices is an interesting but challenging matrix for biomarkers. Blood has a dynamic proteome which varies within an individual depending on time of day [15, 16] and is impacted by the collection protocol [17, 18] and various other factors [19]. Blood is also highly heterogeneous between individuals and analysis of thousands of samples is required to appropriately capture the biological diversity of populations [10, 17, 20]. The steep dynamic range of the blood proteome is an added strain, where highly abundant species like hemoglobin, albumin, and immunoglobulins mask the presence of less abundant proteins. It is expected that the coverage of blood or blood plasma proteome will be limited [21-23], thus there is an impetus to reproducibly quantify a limited set of proteins rapidly and reproducibly across a vast number of samples. The presented platform reproducibly quantified 318 proteins across 87 remote sampling devices from different donors.

Cell based perturbation experiments are another application which requires high throughput. In these studies, cells are grown in individual wells and subjected to various treatments, or combinations of treatments. The effects of these perturbations are then examined by profiling the cell’s molecular species[24-26]. The number of samples in such studies grows exponentially when multiple biological replicates are subjected to several combinations of conditions within multiple time points and dosages. With cell culture automation it is possible to grow multiples of 96-well plates while applying different perturbations across each plate. A reasonable pace of proteomic analysis to match this throughput is one 96-well plate per day per instrument, or 15 minutes per sample. Ideally a high number of proteins representative of most biological pathways can be reproducibly quantified at this pace. With the presented platform over 4,000 proteins were reproducibly quantified in the lysate of AC16 cells.

Two mass spectrometry innovations address the challenge of maximizing the number of quantified analytes within a short analysis time: additional dimension of analyte separation through ion mobility, and rapid high resolution mass analysis. Ion mobility separation obviates the need for long chromatographic separation by providing an orthogonal second dimension that multiplicatively increases peak capacity, which correlates with detectable analytes [27]. A fast mass analyzer is then required to carry out the necessary selective fragmentations for identification and specific quantification [28]. With these innovations, the role of LC becomes rapid and reproducible introduction of the separated analytes into the MS. The ratio of time during which analytes are detected to total time of analysis (instrument utilization, IU) is a critical metric of throughput because it roughly correlates to the number of quantified analytes. However, achieving a high IU during rapid analysis is challenging because the required steps during which useful data is not generated including the auto-sampler injection of the sample, the dwell time during which the analytes traverse the analytical column, and the cleaning and equilibration steps cannot be accelerated with the same flexibility as the analytical gradient.

The presented dual-trap single column configuration is an accessible, easy to implement innovation which increases IU at any throughput, but especially in rapid analysis. This configuration is suitable for high flow rates, micro LC flow rates and nano-flow. In this configuration, one trap is back-flushed with a high organic buffer, equilibrated, and loaded with the subsequent sample while, in parallel, a sample loaded on the other trap is separated on the analytical column and analyzed. With this parallelization a maximum ratio of instrument time is devoted to collection of useful data (**Fig. 1**). Here we present the adaptation of this configuration for analysis of blood from remote sampling devices and cell lysates at 15 minutes per sample throughput. The presented configuration runs at the 9.5 µL/min flowrate and achieves optimal performance with 1,000 ng of peptides injected on column, thus making it suitable for some sample limited applications.

**Figure 1.**
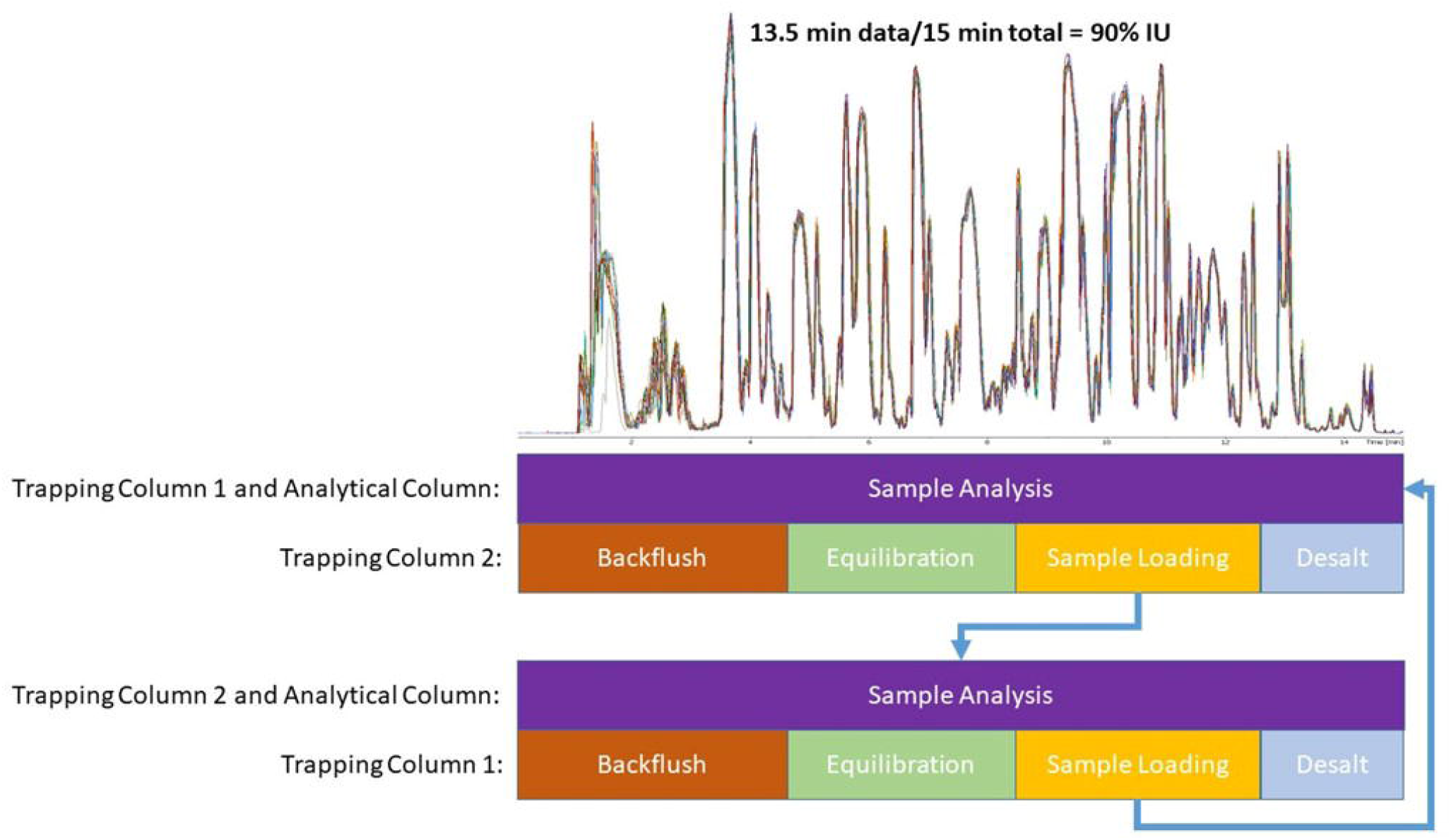
Parallelization of sample analysis on one trap while the second trap is back-flushed, equilibrated, loaded with the next sample, and desalted. The top is an overlay of 10 injections (5 on each trap) demonstrating the reproducibility of the set-up. The dwell time is approximately 1 minute, while 30 seconds are used to flush the analytical column, thus generating 13.5 minutes of data duringa 15 minute run time resulting in 90% instrument utilization (IU).

## Materials and Methods

### LC configuration

The dual-trap single column configuration schematic is illustrated in **Figure 2** and requires a 10-port 2-position valve, a second 2-position valve with at least 6 ports, an analytical pump, and a loading pump. In these experiments an Ultimate 3000 nano-RSLC instrument (Thermo) was used, but any instrument with the aforementioned features can be configured as such. To operate this configuration two complementary instrument methods are programmed. In the first method (**Fig. 2A and B**) the analytical gradient is delivered through Trapping Column 1 and Analytical Column, while the loading pump connected through the auto-sampler delivers a plug of highly organic solvent to clean and back-flush (in reverse direction of the analytical gradient) Trapping Column 2 (**Fig. 2A**). The direction of loading pump flow is switched so that the subsequent sample picked up by the auto-sampler is loaded onto Trapping Column 2 in the analytical direction (forward loaded, **Fig. 2B**) and desalted for several minutes. In the next run the second method reverses the roles of the traps (**Fig. 2C and D**). The methods are alternated for the entire sample set. A blank is analyzed in the first run since no sample was loaded in the previous run and buffer is injected in the final run since there is no subsequent sample. It is important to note that the loading pump is not intended for gradient delivery and while it can be used to deliver a slug of organic buffer and equilibrate back to aqueous composition before sample loading at the 60 µL/min flowrate, it is not able to do so at the nano-liter/min flowrate, so instead the auto-sampler sample loop can be used to inject highly organic buffer during trap column cleaning.

**Figure 2.**
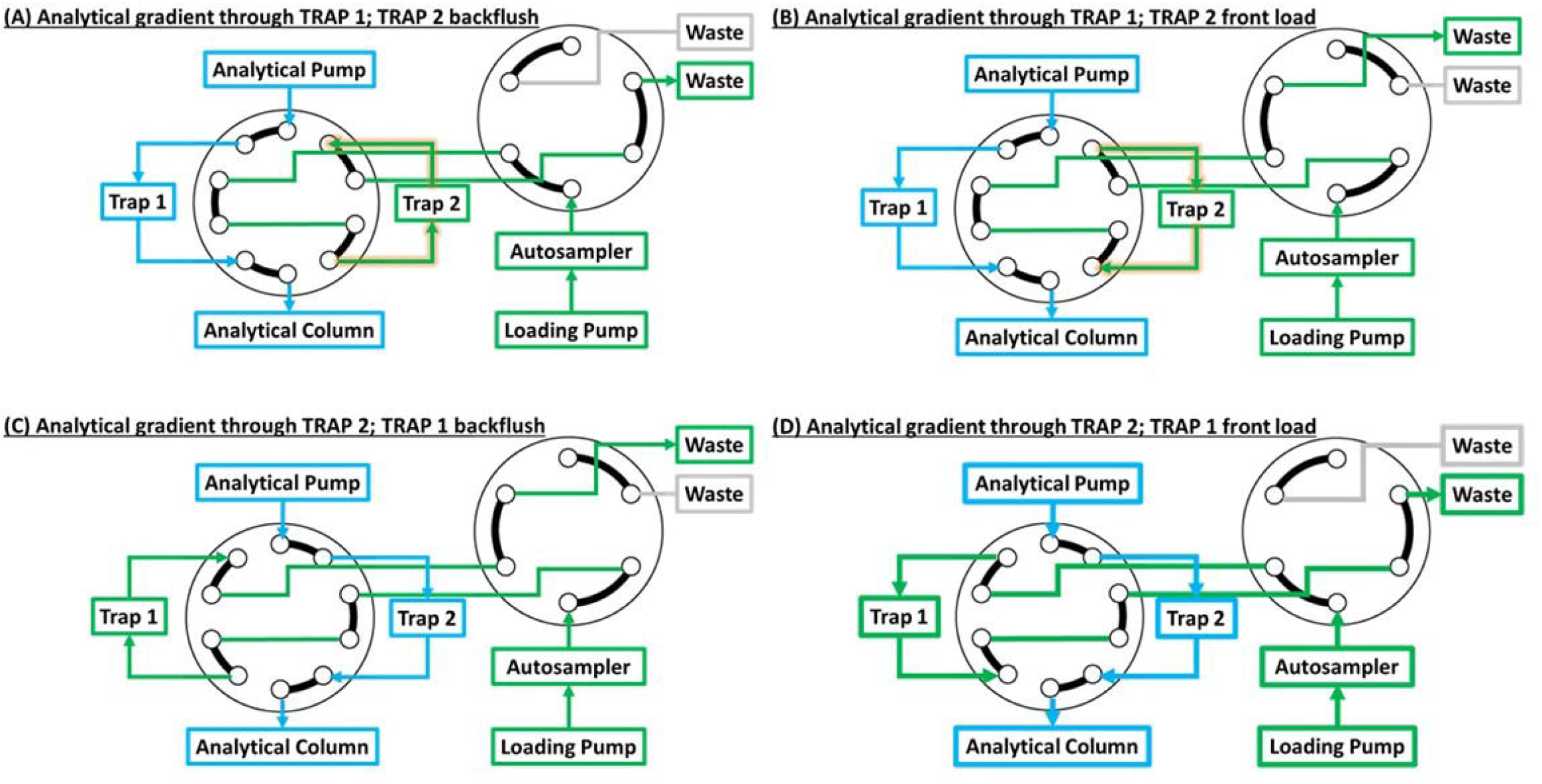
Valve schematic of the dual-trap one-column configuration. The 10 port valve dictates which trap is connected to the analytical pump and column while the other trap is connected to the loading pump and auto-sampler. While the sample loaded on Trap 1 is eluted to the analytical column and analyzed Trap 2 is back-flushed with high organic buffer (Panel A). After equilibration back to aqueous composition the next sample can be loaded in the same direction as the analytical separation (Panel B). This prevents any precipitated contaminants from getting to the analytical column and reduces downtime due to clogging. I n the next run the roles of the traps are reversed (Panels C and D).

### LC-MS Analysis

All samples were analyzed on an Ultimate 3000 nano-RSLC equipped with the capillary flow selector at 9.5 µL/min analytical flowrate. Peptides were trapped on Phenomenex 0.3 mm internal diameter, 150 mm long columns packed with 5 um Kinetex C8 beads, and separated on a CapLC 50 cm micro-pillar array column from Pharmafluidics (now Thermo). An 8-nozzle M3 emitter installed in a MnESI source from Newomics was used to split the flow just prior to electrospray [29, 30]. The entire assembly was connected with 50 um internal diameter Viper capillaries (Thermo) and kept at 50° C in the column oven. The analytical gradient used 0.1% formic acid in water (mobile phase A) and 0.1% formic acid in acetonitrile (mobile phase B) and was delivered as follows: start at 7% B; linear ramp to 22%B over 8.5 minutes; linear ramp to 38%B in 4.3 minutes; jump to 98%B over 0.2 minutes, 0.9 min hold at 98% B, drop back to 7% B and hold for 1 minute (15 minutes total). The loading pump was connected to three solvents: 0.1% formic acid in water (loading buffer A), 0.2% formic acid in 70% acetonitrile 30% water with 5 mM ammonium formate (loading buffer B), 0.2% formic acid in 90% isopropanol with 10% acetonitrile and 5 mM ammonium formate (loading buffer C). Although one highly organic solvent to flush the trapping columns and one aqueous buffer to load the samples is sufficient, this set of buffers was used for easy switching to a different application. At the beginning of the method the loading pump delivered 50% B and 50% C at 60 µL/min for 0.5 minutes while reducing the flowrate to 55 µL/min. The solvent was switched to 100% A over 0.3 minutes and held for 8.2 minutes at 55 µL/min to equilibrate back to aqueous composition. The loading pump flowrate was reduced to 20 µL/min at 9.5 minutes and held for the remainder of the method for sample loading and desalting. The auto-sampler was operated with a user program in which the first command initialized the run, next the auto-sampler filled the 20 µL sample loop with acetonitrile and injected it to help flush the trapping column. Two 25 µL rinses with water removed any organic from the needle assembly. After a 3-minute pause, 20 µL of sample was collected into the loop and injected onto the trapping column, this coincided with the drop of loading pump flowrate to 20 µL/min at 9.5 minute run time. Finally, the syringe was emptied and the needle was washed. The auto-sampler program is presented in **Supplement Table 1**.

Data were acquired on a Bruker TIMS-TOF Pro mass spectrometer using PASEF-DIA. Ions were accumulated for 70 ms, separated with 70 ms ramp, and fragmented in 40 m/z windows which covered the 360 to 1120 m/z and 0.65 to 1.41 1/K0 ion mobility ranges corresponding to the majority of observed multi-charged peptide ions. Each ramp cycle contained 1 to 3 DIA scans resulting in a 0.76 s total cycle time. The DIA isolation scheme is presented in **Supplement Table 2**. Electrospray ionization was performed using a Bruker MnESI source with the 8 nozzle M3 emitter from Newomics with the capillary voltage set to 4800 V, the endplate offset voltage set to 500 V, the nebulizer set to 2.9 Bar, and the Dry Gas set to 6.0 L/min and 200 C. The mass spectrometer and Ultimate 3000 were controlled through Hystar 6.0 with the SII plug-in. The two LC methods and the PASEF-DIA method files are uploaded to LCMSMethods.com under dx.doi.org/10.17504/protocols.io.5qpvob27dl4o/v1.

### Data Analysis

Data were analyzed in DIA-NN 1.8 [31] using the library-free search feature or with libraries generated by gas phase fractionation (GPF). To generate the GPF libraries, pooled samples were analyzed repeatedly using the aforementioned LC settings but with data-dependent acquisition (DDA) to acquire fragmentation spectra for precursors within a restricted mass to charge range (e.g. 300 to 500 m/z) **(Supplement Figure 1>)**. Different precursor m/z ranges were analyzed in each run to span the 300 to 1100 range and one full range experiment was carried out as the reference for retention time alignment during library assembly. The GPF data were analyzed in FragPipe 17.1 [32]. In the first search, the open search pipeline with strict tryptic specificity was used to identify the prevalent post-translational modifications (PTMs) and the detectable protein population. The second search was restricted to the identified proteins and most prevalent PTMs, but with semi-tryptic protein specificity and up to 2 allowed missed cleavages. The peptides identified in the second search at <1% FDR were compiled into a spectral library with EasyPQP. A quick library was generated for DIA method optimization using dried blood spots with just nine DDA runs. A more thorough library was generated for the remote sampling devices by collecting each GPF range in duplicate for each of the four cohorts (64 with an additional alignment reference run). Cell lysates were analyzed using the DIA-NN library-free feature in which the human SwissProt protein database was translated into in silico generated spectra with retention time and ion mobility predicted by machine learning. In all analysis, match between runs (MBR) and double pass mode were enabled and MS1 and MS2 mass errors were set to 15 ppm.

For cohort comparison of Mitra device blood the DIA-NN reported protein intensities were log base 2 transformed and standardized across each sample so that the distribution of protein quantities in each sample had a mean of zero and standard deviation of one (**Fig. 5**). UMAP (Uniform Manifold Approximation and Projection for Dimension [33] was used to project the protein quantities onto two dimensions to allow for visualization of protein clustering by cohort. Proteins quantified in every sample (318 proteins) were compared between the control group and the three hypertension cohorts as a combined second group using t-tests with Benjamini-Hochberg correction. The proteins with significantly different means (adjusted p-values < 0.05) were used as inputs to train one of five classification models (Gradient Boosting Decision Trees [GB], Support Vector Classification [SVC], Random Forest [RF], Extra-Trees [ET], and Logistic Regression [LR]) [34-38] using 70% of the data. The remaining 30% of the data was used as the final test set and the model with the highest precision recall area under the curve (PR AUC) was selected as the classifier for hypertension prediction. This data normalization, analysis, and model training were performed in Python version 3.7.11.

### Dried blood spots (DBS) sample preparation

Pooled, mixed-gender blood sample (Golden West Biosolutions, LLC., Human Whole Blood Lot# PS1000.104) was spotted onto Whatman cards. The discs with dried blood spots were transferred to 96 well deep-well plates (1.1 mL volume, Thermo Fisher Scientific). The proteins were solubilized in lysis buffer composed of 9M Urea, 0.03 M TCEP (Thermo Fisher Scientific), and 0.2 M Trizma (Sigma Aldrich) by shaking at 1200 rpm for 60 minutes at 37C. The remainder of the protocol was performed automatically on a Beckman i7 automated workstation (Beckman Coulter). The samples were alkylated in 0.05 M iodoacetamide (IAA). Then the samples were diluted with 0.2 M Trizma to a final volume of 567uL and trypsin was added to 0.15 mM final concentration. The plate was incubated at 42° C for 4 hours with slight shaking at 120 rpm and digestion was quenched by acidification with formic acid. Digested samples were mixed with 2% phosphoric acid, 0.1% formic acid and automatically desalted on an Oasis 30 µm HLB 96-well plate (Waters) following the manufacturer’s protocol adapted for the Beckman i7 and the positive pressure apparatus module to push the solvents through [39].

### Mitra device dried blood sample preparation

Blood was obtained from 87 patients using Mitra devices (Neoteryx). The patients were grouped by blood pressure into cohorts: Cohort 1 did not have high blood pressure (n=37), Cohort 2 had blood pressure between 120/80 and 130/90 (n=29), Cohort 3 had a BP above 130/90 (n=11) and Cohort 4 had high blood pressure but was managing it with medication (n=10). The protein content was extracted from the Mitra devices by 1-hour incubation at 60° C in 35% TFE buffer containing 40 mM dithiothreitol (DTT; Sigma) and 50 mM ammonium bicarbonate (Sigma). The rest of the sample processing was performed automatically on the Beckman i7. Iodoacetamide was added to 10 mM concentration and the samples were incubated at 25° C in the dark for 30 minutes. An addition of 5 mM DTT quenched alkylation and addition of 50 mM NH4CO3 diluted TFE to 5% final concentration. Trypsin was added at a ratio of 1:25 protease to substrate and samples were digested for 4 hours at 43° C. Digestion was quenched with addition of formic acid to 1% final concentration. Samples were diluted to 50 ng/ µL concentration before LC-MS analysis [40].

### Cell Lysate Preparation

The AC16 cell line was grown in house in high glucose Dulbecco’s modified eagle medium: nutrient mixture F-12 (DMEM/F12; Gibco), with 12.5 % fetal bovine serum (FBS; Gibco) supplemented with antibiotic-antimycotic (100X; Gibco) at 37 °C (5% CO2). At 1.44 × 10^7^ cells/mL concentration the cells were washed two times with PBS, trypsinized, and pelleted at 180 x g. The pellet was washed twice and aliquots of 3.7 × 10^4^ cells suspended in 20 µL of 100 mM ammonium bicarbonate (MilliporeSigma) buffer were loaded into individual wells on three 96-well AFA-Tube TPX plates (COVARIS). Cells were sonicated and lysed for 5 minutes using the LE220-Plus focused-ultrasonicator (COVARIS) using 350 power peak intensity (PIP), 25 % duty factor (DF) and 200 cycles per burst (CPB). Each sample was reduced with 20 mM dithiothreitol (DTT; Pierce) for 10 minutes at 60° C, and then alkylated with 40 mM Iodoacetamide (IAA; BioUltra) for 30 minutes at room temperature in dark. The sample volume was adjusted to 150 uL and to 10% acetonitrile. The samples were digested with sequencing grade modified trypsin from Promega at a ratio of 1:20 (w/w) for 2 hours at 42°C. The samples were acidified to 0.6% Trifluoroacetic acid Optima LC/MS grade (TFA; Fisher Scientific) and dried under vacuum. Prior to analysis the dried peptides were reconstituted in 80 µL of 0.1% formic acid, to load 1,000 ng in 20 µL injections.

## Results and Discussion

With the dual-trap one-column configuration one trapping column is cleaned and loaded with the following sample during analysis of the sample loaded on the second trap. A typical auto-sampler injection requires 2.5 minutes and the system cleaning and equilibration is reduced by 0.5 minutes, thus saving 3 minutes per run which amounts to a 20% increase in throughput at 15 minutes per sample. Parallelization has previously been implemented using configurations where two alternating separations are performed with two analytical columns driven by two analytical pumps with and without corresponding trapping columns [41, 42]. The presented configuration has several advantages over these more complex set-ups. First, LCs equipped with a loading pump and a binary analytical pump (e.g. Ultimate 3000 nano-RSLC and Waters Acuity) are widely implemented making this strategy accessible to many researchers. The dual column strategy can use a valve to select the column that is connected to the electrospray source during analyte elution[43], however this post-column volume reduces performance at the nano- and low micro-liter/min flowrates and a custom electrospray source that can accommodate two separate emitters is required [44]. Furthermore, a second analytical column is an additional source of variability. The presented approach is robust and reproducible at the low-microliter/min flowrate and lower and works with conventional single emitter sources which makes it suitable for low sample quantity applications.

In the presented implementation of this configuration, peptides were quantified for 13.5 minutes out of the 15 minute run time thus achieving a 90% IU. The DIA data acquisition scheme was optimized using a standard composed of digested dried blood spots and the isolation windows were selected to cover the majority of detectable multi-charged peptide ions. Data acquisition schemes with varying ramp times and isolation window widths were evaluated by assessing the peptide level quantitative reproducibility of 10 injections of 1,000 ng of the standard (**Supplement Figure 1**). The highest number of identified precursors (3,985 total, 3,697 on average) and the lowest CV (23% median, 26% average) correlated with the highest number points across the elution of each precursor, indicated by the 3.1 MS2 scans across the full width half max (FWHM). To further exploit this trend, the isolation windows were optimized to increase the MS2 scans at FWMH and the method described in the Methods section averaged 3.3 to 4 MS2 scans at FWHM and was used for the subsequent experiments.

A loading curve spanning 10 to 1,500 ng of the DBS standard was generated to establish optimal loading quantity and evaluate quantitative variance at each level. Each quantity was injected 4 times, except for the 1,000 ng and 1,500 ng levels which were injected 10 times to serve as the first day reproducibility set. One of the 25 ng injections was not picked up by the auto-sampler and was excluded from further analysis. The number of precursor and protein identifications and the %CV at each level is presented in **Figure 3A and B**. Based on this data the highest, most reproducible identifications were achieved at 1,000 ng. Thus the intra and inter day reproducibility was tested with ten 1,000 ng injections on two additional days and the %CV distribution is presented in **Figure 3C and D**. On two of the days 128 and 134 proteins had a CV of less than 10%, but on the second day, which reduced the inter-day reproducibility overall only 96 proteins were under this threshold. Despite this, 234 proteins had a CV below 20% on all three days and 212 met this threshold inter-day. If an inter-day variance of 30% CV is acceptable, then 291 proteins and 2,999 precursors (or 4,177 at 40% CV) can be considered reproducibly quantifiable with this platform.

Next, we applied the platform to real-world blood samples collected using Mitra devices. The specimens were grouped by blood pressure (BP): Cohort 1 had normal BP (n= 37), Cohort 2 had BP between 120/80 and 130/90 (n=29), Cohort 3 had a BP above 130/90 (n=11) and Cohort 4 had high BP that was treated with medication (n=10). On average, 366 proteins were quantified in each cohort with 318 proteins quantified in every sample and 362 proteins quantified in 80% of the samples (**Fig 4A, 4B**). At the precursor level, 2,166 were quantified in every sample, while 4483 were quantified in 80% of the samples with a total of 5,500 to 6,000 precursors quantified in most samples (**Fig. 4**). The reproducibility was excellent based on uncorrected log2 transformed protein quantity distributions across all 87 samples (**Fig 5A**), and standardization neatly aligned the quantity distributions (**Fig 5B**). UMAP was used for dimension reduction to examine the unbiased relationship between the 4 cohorts (**Fig 5C**). Remarkably, cohorts 3 and 4 cleanly separated from the intermixed control and lower hypertension cohorts along the second dimension. To separate the control cohort from a second group of the three hypertension cohorts, machine learning classification models were trained based on proteins which were quantified in every sample and significantly differentiated between cohort 1 and the combined hypertension cohorts. Data was split into 70% training and 30% test sets, and various models were optimized using random hyperparameter searches before assessing performance on the test set. Of all the models, extra-trees had the highest PR AUC score of 0.91 (**Supplement Table 3**), which was substantially better than the no-skill model (**Fig 5D**). Altogether, this analysis shows that data generated from difficult samples using this high throughput platform can be used to distinguish patient groups even when the underlying biology is subtle.

**Figure 3.**
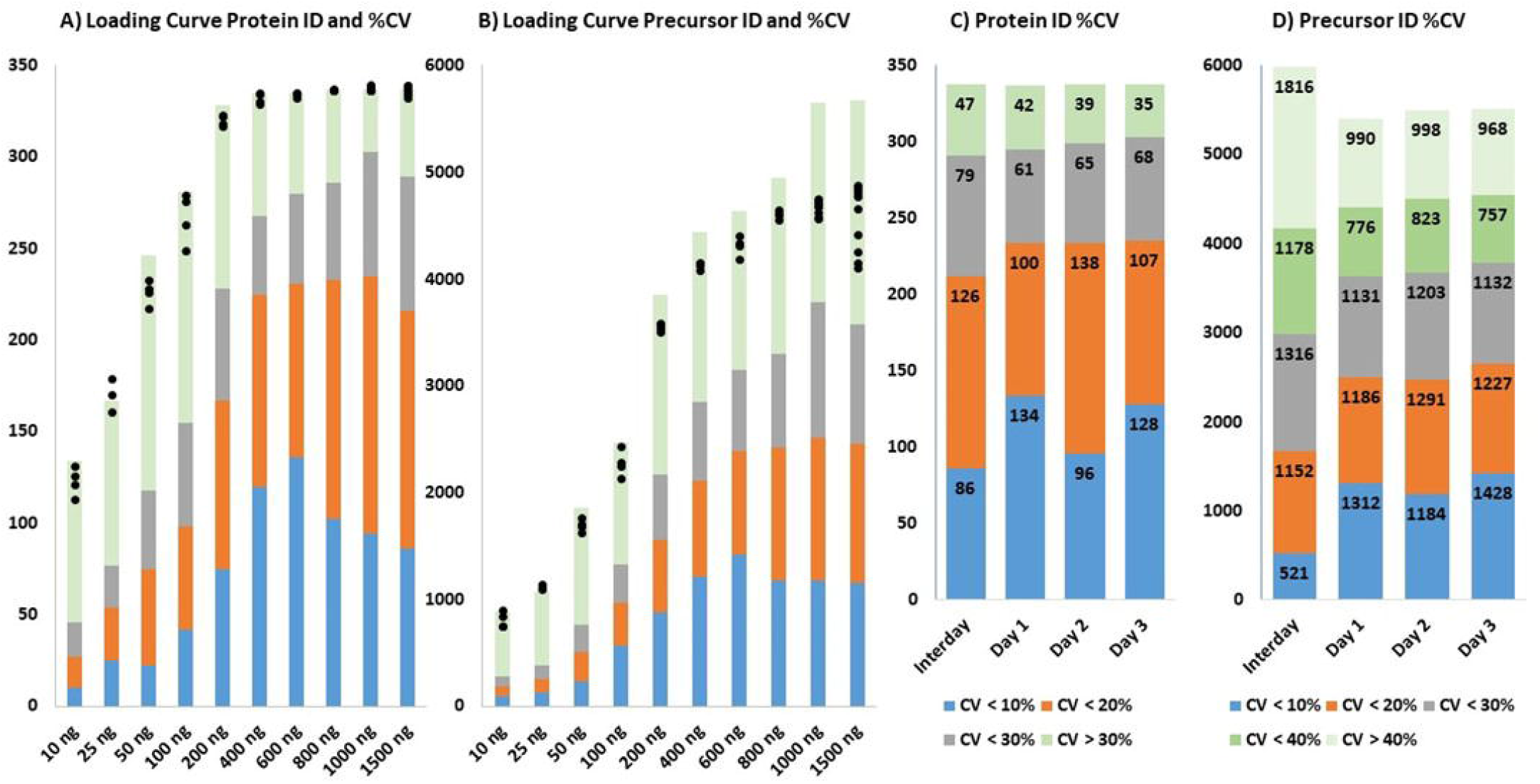
Loading curve and Reproducibility. Panels A and B represent the protein and precursor ion identifications respectively at each amount of dried blood spot peptides. The black points represent the number of identifications in each replicate of the analysis. The colors represent the number of proteins or precursors meeting the %CV criteria, blue is CV under 10%, orange is CV10-20%, grey is CV 20-30%, and green is CV over 30%. Panels C and D represent the protein and precursor ion reproducibility on three different days based on 10 replicate injections and the inter-day reproducibility. The bars represent the number of proteins and precursors meeting the CV criteria. The same color code is repeated from Panels A and B, except for precursor level reproducibility the categories of 30-40% (green) and over 40% (paler green) are added.

**Figure 4.**
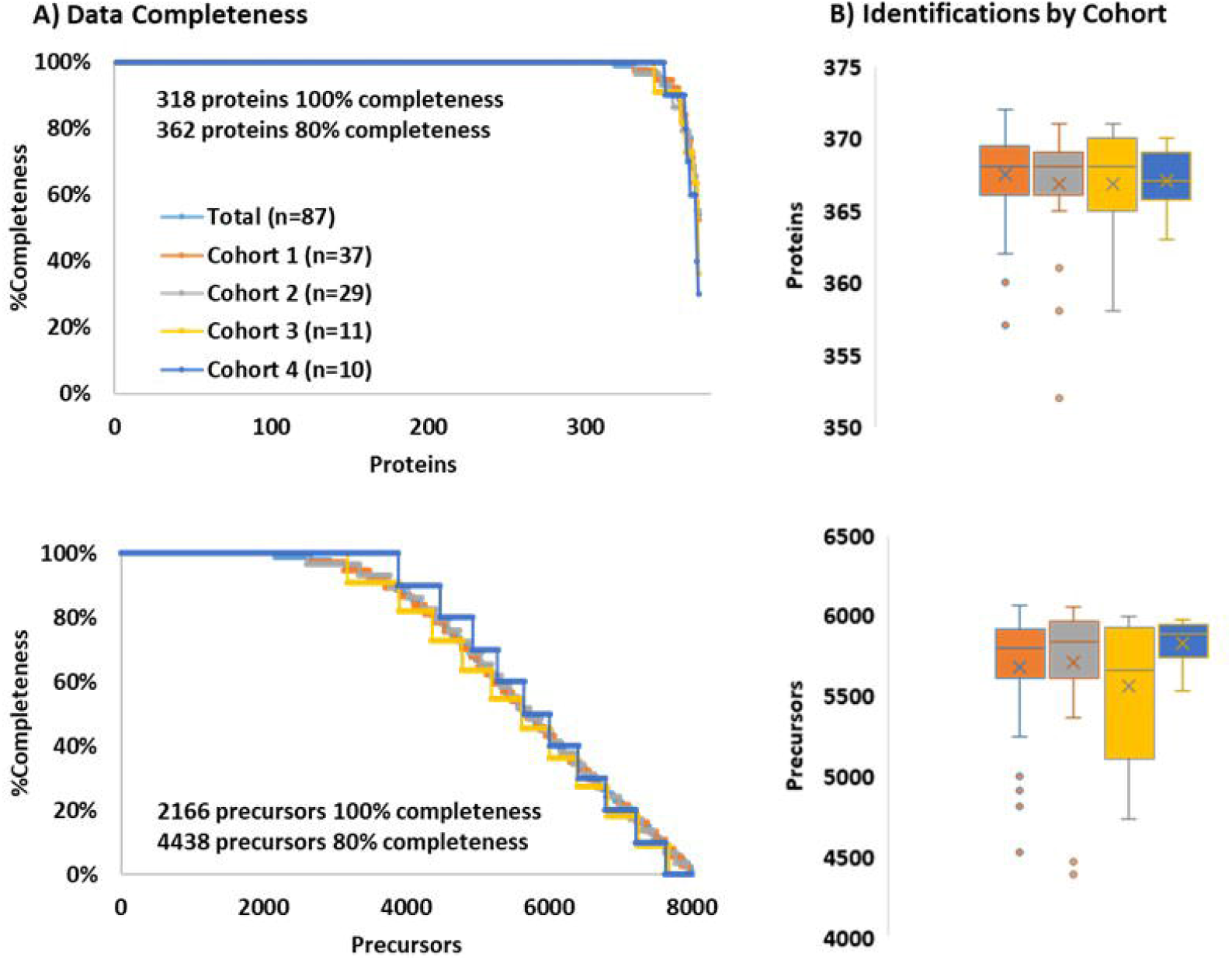
Data completeness (A) and identifications (B) by cohorts in Mitra remote sampling device extracted blood samples. Panel A shows the total data completeness at the protein (top) and precursor ion (bottom) level. The curves for total (light blue), Cohort 1 (orange), Cohort 2 (grey), Cohort 3 (yellow). Cohort 4 (dark blue), are overlaid, at both data levels. Panel shows the identification distribution at the protein (top) and precursor ion (bottom) levels for each cohort following the color scheme from Panel A.

**Figure 5.**
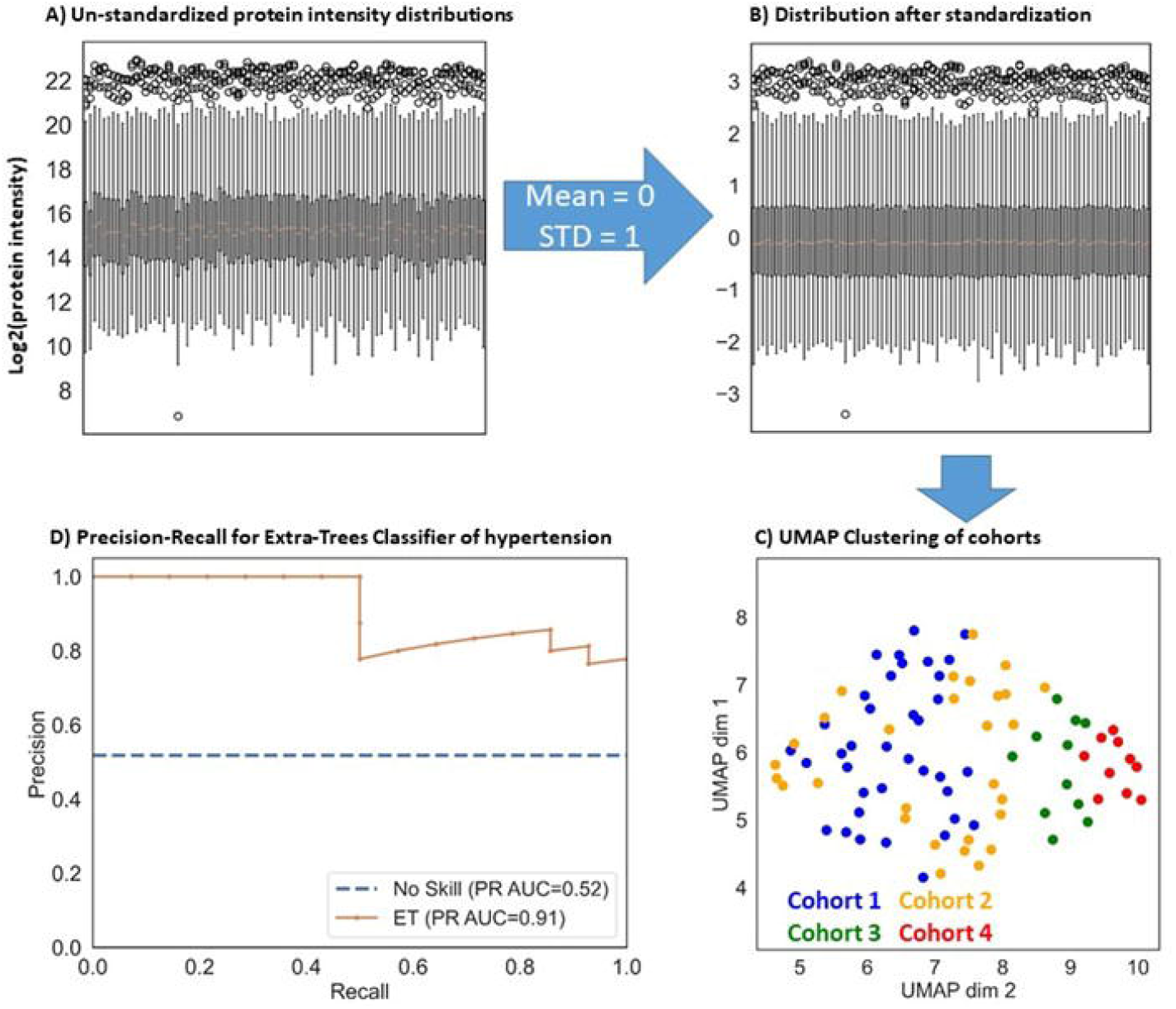
Discrimination of hypertension cohorts based on high throughput analysis of remote device blood samples. Panels A and B show the standardization of protein intensities to enable UMAP clustering (Panel C) of the different cohorts. Cohorts 3 and 4 cleanly separate along the second dimension. While a subset of Cohort 2 separates from Cohort 1 on the second UMAP dimension the two groups are intertwined. However a classifier was trained to distinguish Cohort 1 (the control group) from the remaining hypertension cohorts (Panel D).

The platform was evaluated for rapid analysis of cell lysates in perturbation studies. Three identical 96-well plates containing untreated AC16 cell lysate were prepared. This experiment evaluated the depth of analysis which can be accomplished in 15 minutes and the combined reproducibility of the sample preparation protocol and analytical set-up. The acquired data were analyzed against the human SwissProt database using the library-free search option of DIA-NN. Out of 296 samples, 9 injections failed because the auto-sampler did not pick up the sample, establishing a 3% re-injection rate that can be improved with a more robust auto-sampler. On average, 37.5 k precursor ions and 4,534 proteins were identified across the 279 successful injections. The distribution of identifications by plate is presented in **Figure 6A**. The dataset completeness and reproducibility both at the protein and precursor level are presented in **Figure 6B**; 13,165 precursors and 3,515 proteins were identified in all 279 samples, if a rate of 5% missing values is acceptable then these numbers increase to 22,621 and 3,977 for precursors and proteins, respectively. For the core proteins and precursors (those detected in all successful injections), the average %CV was 11% and 19%, respectively.

**Figure 6.**
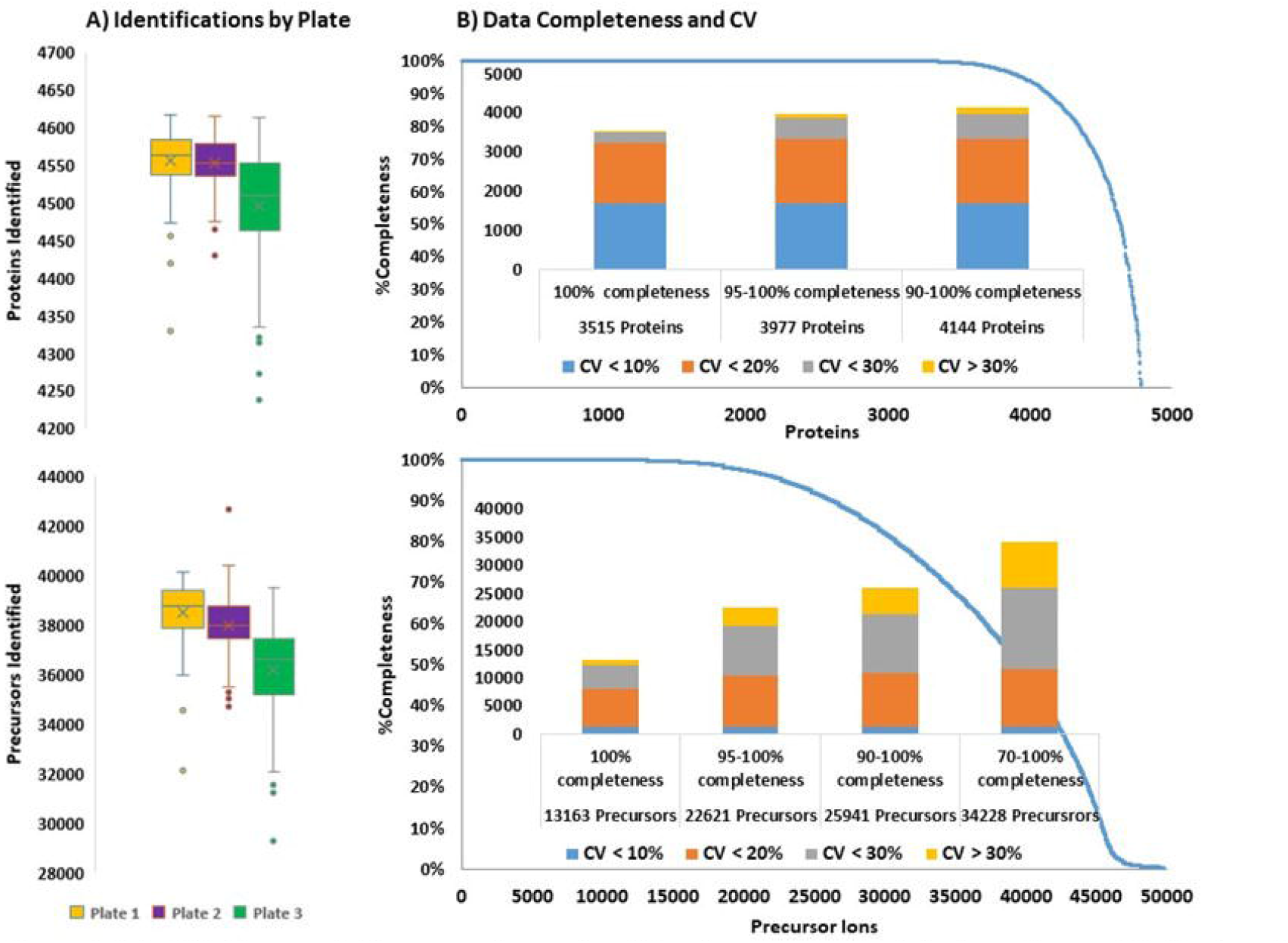
Identifications and data completeness in cell lysate samples. Panel A shows the protein (top) and precursor ion (bottom) identification distribution on each plate. Panel 6 shows the data completeness curve for each protein (top) precursor (bottom) over all samples. The bar graph represents the %CV distributions at each data completeness level: blue CV < 10%; orange CV 10-20%; grey CV 20-30%; and yellow CV over 30%.

The presented assembly robustly and rapidly analyzes challenging physiological samples like blood and achieves deep profiling in cell lysates. A key consideration for developing this platform is the ability to perform with minimal down time. Thus fairly large traps were selected in the expectation that they would not clog and will be able to filter out any contaminants that may compromise analysis. Another key consideration is variability in performance between the two traps. We address this potential issue by routinely running trap synchronization QC, in which the base-peak chromatograms from several runs on each trap are overlaid to ensure that the retention time and peak widths are identical on both traps. A failure in this alignment indicates that there is possibly a leak in one of the connections, some contamination is influencing trap retention, or the trap needs to be replaced. The analytical flowrate was set at 9.5 µL/min because the Ultimate 3000 operates at a maximum of 10 µL/min with the capillary flow module. The CapLC column was selected because it can robustly process 1 to 3 thousand samples, but it does not tolerate a pressure above 350 bar. At the analytical flowrate the back pressure was 200-250 bar, so it is possible to increase the flowrate by several microliters/min and further reduce the dwell time and increase IU. The IU can also be increased by reducing the dwell volume with smaller trapping columns or a different analytical column, but selection of trapping columns which have identical performance and can operate robustly requires thorough testing. It is important to note that while the focus of this publication was tryptic peptides the described configuration can be used to rapidly profile other sets of analytes such as metabolites, lipids, intact proteins, and RNA with the appropriate chromatographic columns and mobile phases. The raw data presented in this manuscript is shared through the MASSIVE repository (DOI#XXXXX)

## Conclusion

We’ve demonstrated that the dual-trap one-column configuration achieves 90% IU at 15 minute per sample throughput in two key LC-MS applications: analysis of cell lysates in a perturbation study and analysis of clinical specimens collected using remote blood sampling devices. We envision implementing this platform synchronously with 24-hour incubation of cells cultured on 96 well-plates with different conditions applied across the plate. This will allow zero down time in perturbation studies as samples are incubated for 24 hours, digested the next day, and analyzed the following day while the subsequent plates are prepared and digested in parallel. Our platform quantified over 4,000 proteins in a human cardiomyocyte cell line (AC16) lysate, which corresponds to approximately 20% of the total genome in 15 minutes. The platform is also versatile as demonstrated in the analysis of blood from remote sampling devices. In combination with remote sampling, this platform can be used to execute ambitious surveys of large cohorts with multiple time-points. Even at the cursory depth of the top 300-400 blood proteins we were able to distinguish samples based on the blood pressure phenotype using machine learning using data from just 15 minutes of instrument time. Similar algorithms can be generated for all sorts of phenotypes and disease states and greater depth can be achieved through longer methods or fractionation. Our dual-trap single-column platform provides a significant boost in throughput without sacrificing the quality of the generated data thus empowering precise and reproducible quantitation across large samples sets and allowing proteomics to capture population and biological heterogeneity.

## Acknowledgements

We would like to thank our collaborators at Bruker for instrument support especially Michael Krawitzky, Francesco Pingitore, and Christopher Adams. We would also like to thank our collaborators from Newomics for help in optimization of electrospray ionization settings especially Daojing Wang and Mark J. Schroeder.

