## Supplement Figures and Tables for "Parallelization with Dual-Trap Single-Column Configuration Maximizes Throughput of Proteomic Analysis"

### Slide 1
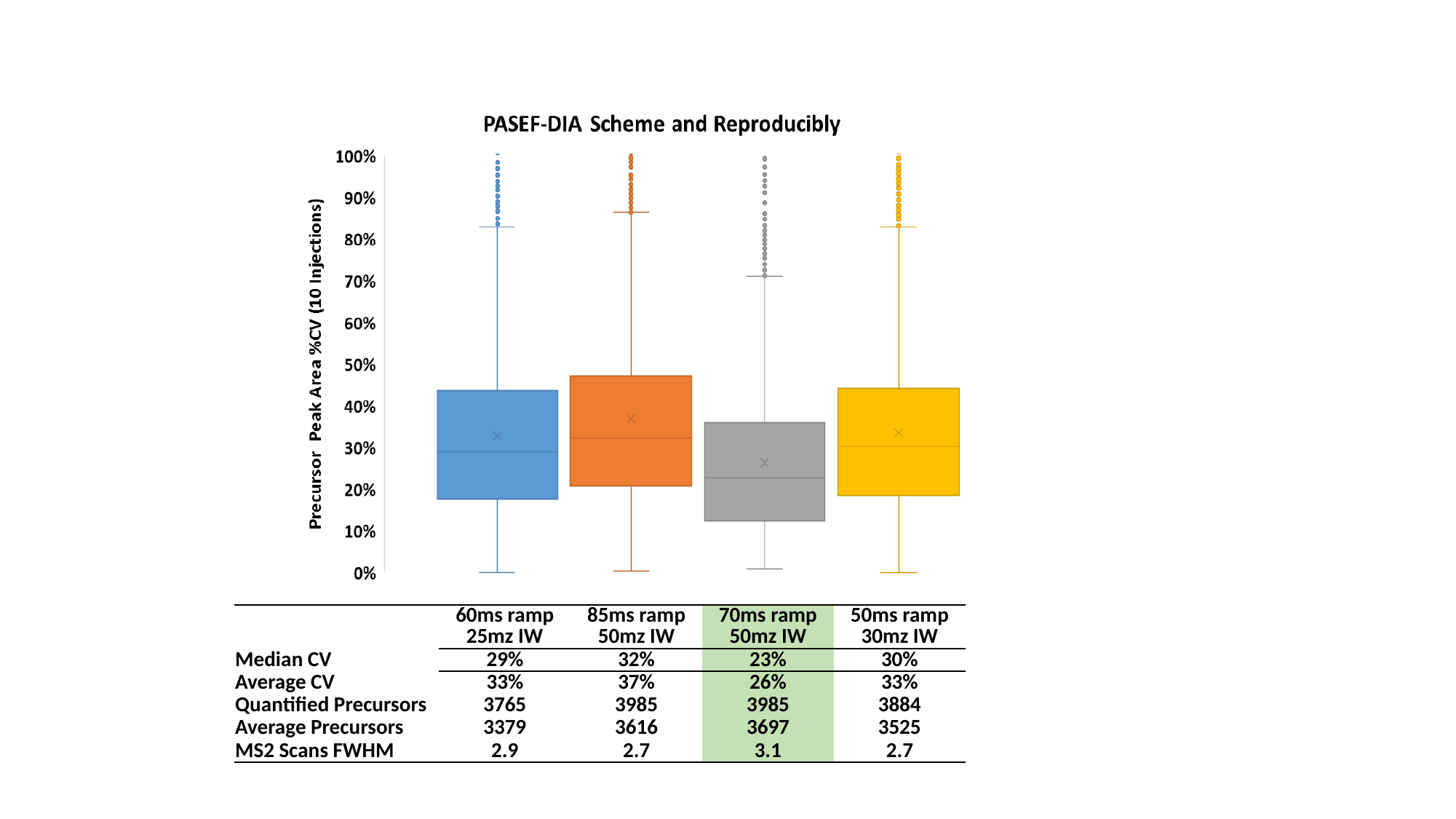

| | 60ms ramp 25mz IW | 85ms ramp 50mz IW | 70ms ramp 50mz IW | 50ms ramp 30mz IW |
| --- | --- | --- | --- | --- |
| Median CV | 29% | 32% | 23% | 30% |
| Average CV | 33% | 37% | 26% | 33% |
| Quantified Precursors | 3765 | 3985 | 3985 | 3884 |
| Average Precursors | 3379 | 3616 | 3697 | 3525 |
| MS2 Scans FWHM | 2.9 | 2.7 | 3.1 | 2.7 |

### Slide 2
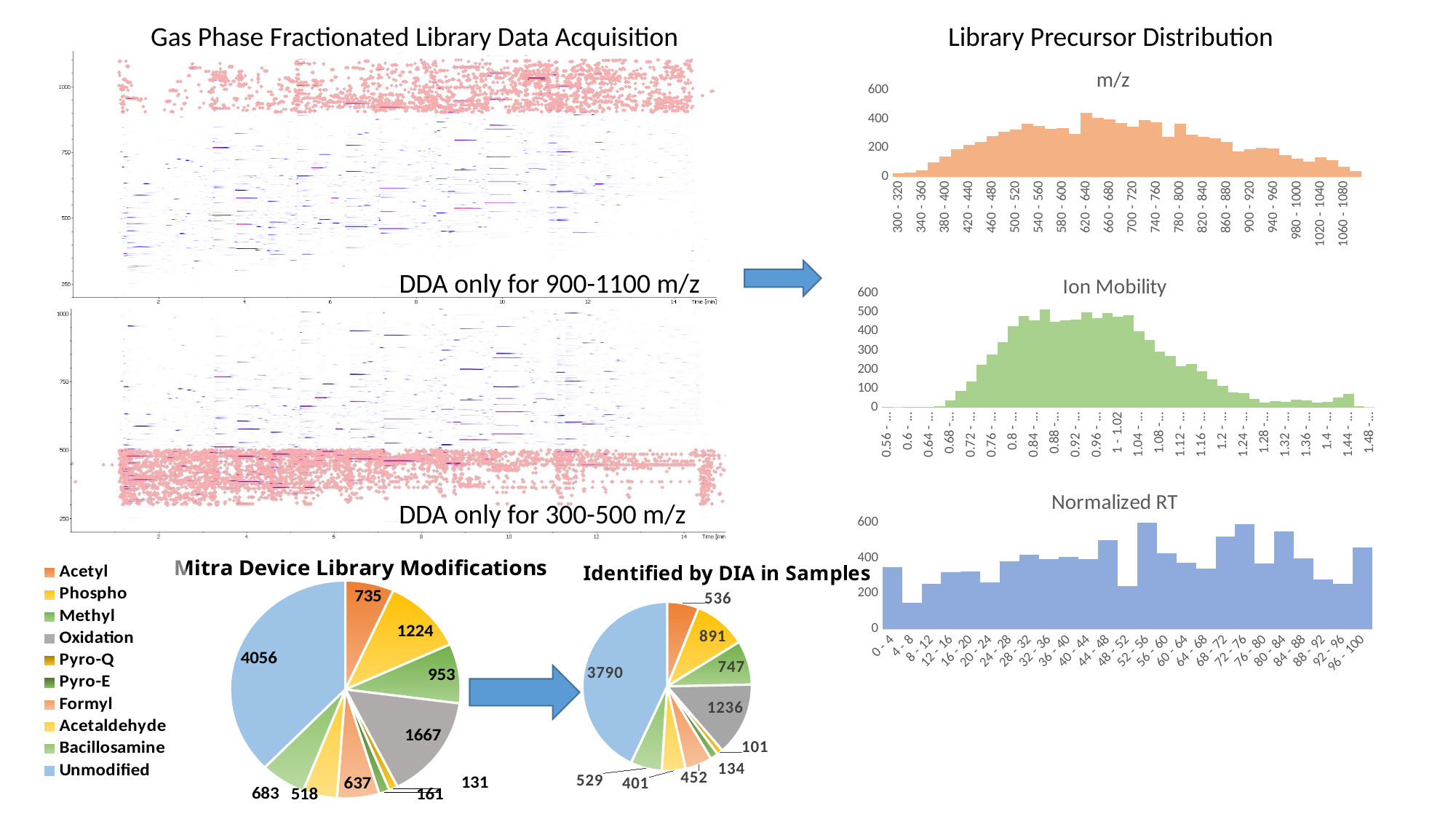

Gas Phase Fractionated Library Data Acquisition
Library Precursor Distribution
#### Chart:
| Category | m/z |
|---|---|
| 300 - 320 | 24.0 |
| 320 - 340 | 27.0 |
| 340 - 360 | 44.0 |
| 360 - 380 | 101.0 |
| 380 - 400 | 139.0 |
| 400 - 420 | 193.0 |
| 420 - 440 | 223.0 |
| 440 - 460 | 240.0 |
| 460 - 480 | 284.0 |
| 480 - 500 | 313.0 |
| 500 - 520 | 327.0 |
| 520 - 540 | 367.0 |
| 540 - 560 | 353.0 |
| 560 - 580 | 334.0 |
| 580 - 600 | 336.0 |
| 600 - 620 | 296.0 |
| 620 - 640 | 441.0 |
| 640 - 660 | 410.0 |
| 660 - 680 | 399.0 |
| 680 - 700 | 374.0 |
| 700 - 720 | 349.0 |
| 720 - 740 | 395.0 |
| 740 - 760 | 376.0 |
| 760 - 780 | 275.0 |
| 780 - 800 | 368.0 |
| 800 - 820 | 293.0 |
| 820 - 840 | 279.0 |
| 840 - 860 | 266.0 |
| 860 - 880 | 240.0 |
| 880 - 900 | 177.0 |
| 900 - 920 | 192.0 |
| 920 - 940 | 203.0 |
| 940 - 960 | 195.0 |
| 960 - 980 | 148.0 |
| 980 - 1000 | 123.0 |
| 1000 - 1020 | 105.0 |
| 1020 - 1040 | 136.0 |
| 1040 - 1060 | 114.0 |
| 1060 - 1080 | 71.0 |
| 1080 - 1100 | 41.0 |DDA only for 900-1100 m/z
#### Chart: Ion Mobility
| Category | |
|---|---|
| 0.56 - 0.58 | 4.0 |
| 0.58 - 0.6 | 0.0 |
| 0.6 - 0.62 | 1.0 |
| 0.62 - 0.64 | 1.0 |
| 0.64 - 0.66 | 2.0 |
| 0.66 - 0.68 | 8.0 |
| 0.68 - 0.7 | 36.0 |
| 0.7 - 0.72 | 88.0 |
| 0.72 - 0.74 | 138.0 |
| 0.74 - 0.76 | 225.0 |
| 0.76 - 0.78 | 278.0 |
| 0.78 - 0.8 | 342.0 |
| 0.8 - 0.82 | 428.0 |
| 0.82 - 0.84 | 480.0 |
| 0.84 - 0.86 | 458.0 |
| 0.86 - 0.88 | 516.0 |
| 0.88 - 0.9 | 450.0 |
| 0.9 - 0.92 | 459.0 |
| 0.92 - 0.94 | 461.0 |
| 0.94 - 0.96 | 501.0 |
| 0.96 - 0.98 | 471.0 |
| 0.98 - 1 | 497.0 |
| 1 - 1.02 | 479.0 |
| 1.02 - 1.04 | 486.0 |
| 1.04 - 1.06 | 400.0 |
| 1.06 - 1.08 | 355.0 |
| 1.08 - 1.1 | 295.0 |
| 1.1 - 1.12 | 270.0 |
| 1.12 - 1.14 | 216.0 |
| 1.14 - 1.16 | 227.0 |
| 1.16 - 1.18 | 191.0 |
| 1.18 - 1.2 | 147.0 |
| 1.2 - 1.22 | 113.0 |
| 1.22 - 1.24 | 78.0 |
| 1.24 - 1.26 | 74.0 |
| 1.26 - 1.28 | 46.0 |
| 1.28 - 1.3 | 25.0 |
| 1.3 - 1.32 | 33.0 |
| 1.32 - 1.34 | 30.0 |
| 1.34 - 1.36 | 40.0 |
| 1.36 - 1.38 | 38.0 |
| 1.38 - 1.4 | 24.0 |
| 1.4 - 1.42 | 31.0 |
| 1.42 - 1.44 | 51.0 |
| 1.44 - 1.46 | 71.0 |
| 1.46 - 1.48 | 8.0 |
| 1.48 - 1.5 | 0.0 |
#### Chart: Normalized RT
| Category | |
|---|---|
| 0 - 4 | 347.0 |
| 4 - 8 | 148.0 |
| 8 - 12 | 254.0 |
| 12 - 16 | 318.0 |
| 16 - 20 | 324.0 |
| 20 - 24 | 263.0 |
| 24 - 28 | 381.0 |
| 28 - 32 | 419.0 |
| 32 - 36 | 394.0 |
| 36 - 40 | 405.0 |
| 40 - 44 | 392.0 |
| 44 - 48 | 499.0 |
| 48 - 52 | 241.0 |
| 52 - 56 | 636.0 |
| 56 - 60 | 424.0 |
| 60 - 64 | 371.0 |
| 64 - 68 | 338.0 |
| 68 - 72 | 519.0 |
| 72 - 76 | 590.0 |
| 76 - 80 | 370.0 |
| 80 - 84 | 547.0 |
| 84 - 88 | 398.0 |
| 88 - 92 | 280.0 |
| 92 - 96 | 252.0 |
| 96 - 100 | 460.0 |DDA only for 300-500 m/z
#### Chart: Mitra Device Library Modifications
| Category | |
|---|---|
| Acetyl | 735.0 |
| Phospho | 1224.0 |
| Methyl | 953.0 |
| Oxidation | 1667.0 |
| Pyro-Q | 131.0 |
| Pyro-E | 161.0 |
| Formyl | 637.0 |
| Acetaldehyde | 518.0 |
| Bacillosamine | 683.0 |
| Unmodified | 4056.0 |
#### Chart: Identified by DIA in Samples
| Category | |
|---|---|
| Acetyl | 536.0 |
| Phospho | 891.0 |
| Methyl | 747.0 |
| Oxidation | 1236.0 |
| Pyro-Q | 101.0 |
| Pyro-E | 134.0 |
| Formyl | 452.0 |
| Acetaldehyde | 401.0 |
| Bacillosamine | 529.0 |
| Unmodified | 3790.0 |

### Slide 3
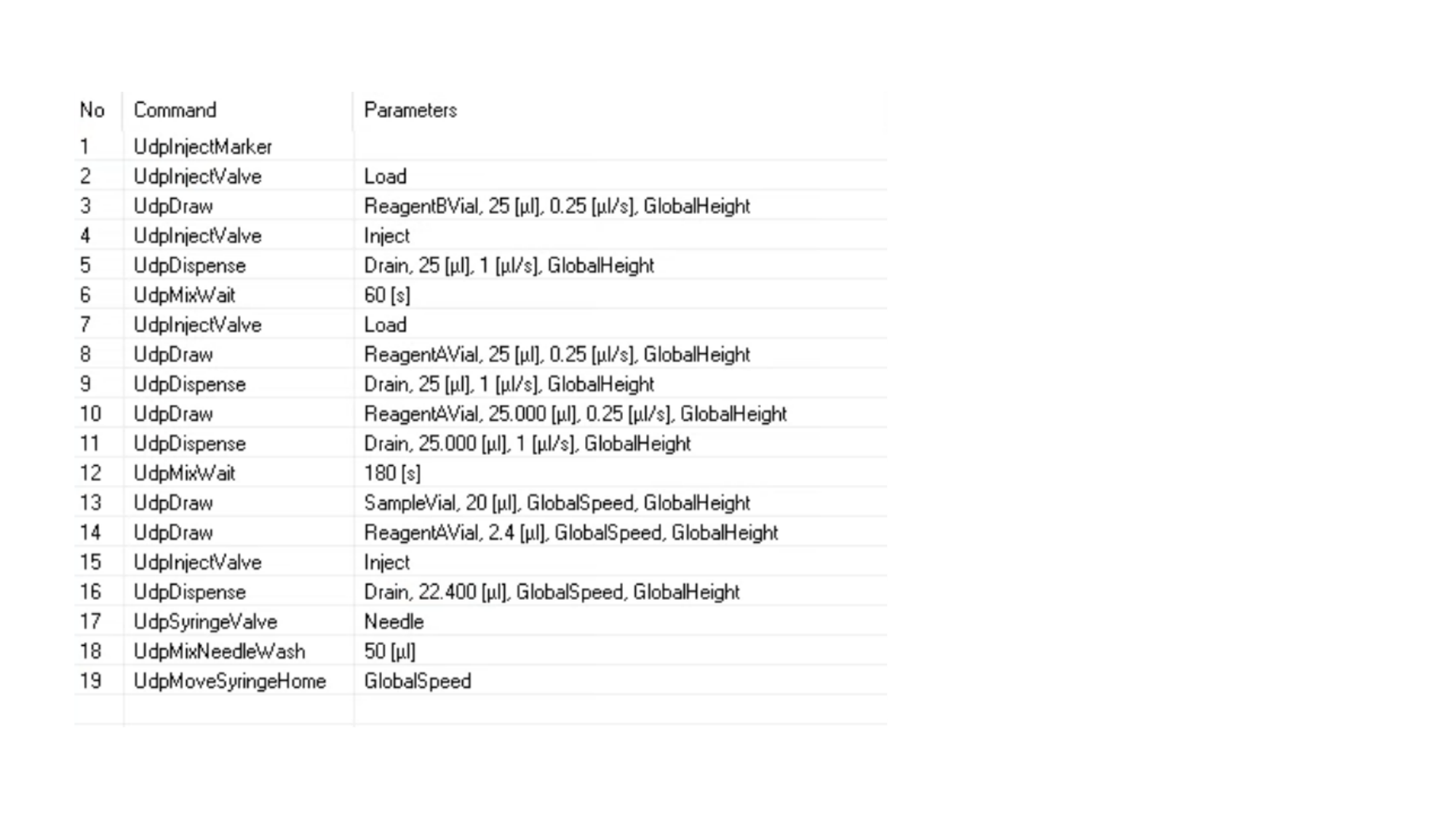

### Slide 4
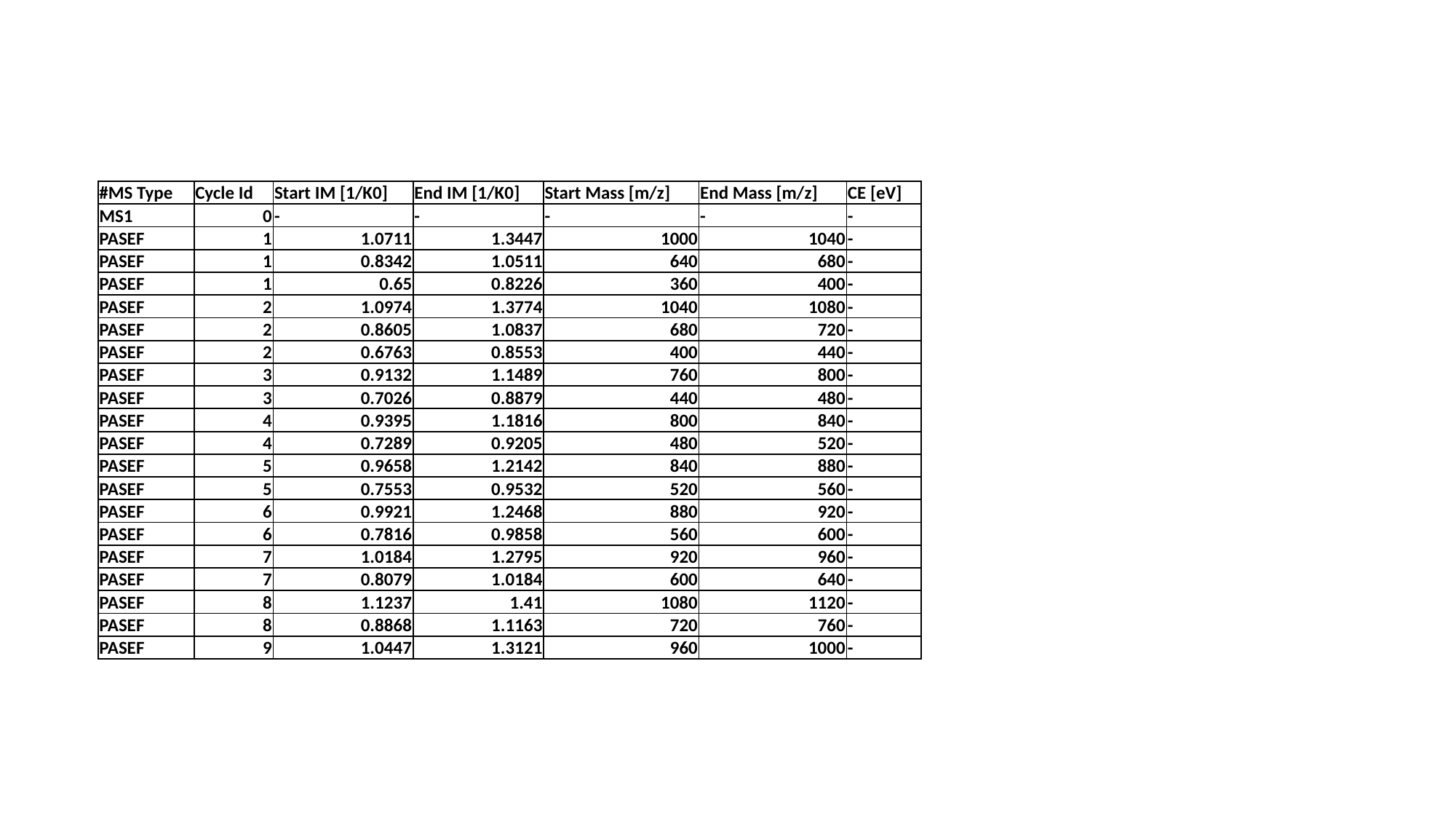

| #MS Type | Cycle Id | Start IM [1/K0] | End IM [1/K0] | Start Mass [m/z] | End Mass [m/z] | CE [eV] |
| --- | --- | --- | --- | --- | --- | --- |
| MS1 | 0 | - | - | - | - | - |
| PASEF | 1 | 1.0711 | 1.3447 | 1000 | 1040 | - |
| PASEF | 1 | 0.8342 | 1.0511 | 640 | 680 | - |
| PASEF | 1 | 0.65 | 0.8226 | 360 | 400 | - |
| PASEF | 2 | 1.0974 | 1.3774 | 1040 | 1080 | - |
| PASEF | 2 | 0.8605 | 1.0837 | 680 | 720 | - |
| PASEF | 2 | 0.6763 | 0.8553 | 400 | 440 | - |
| PASEF | 3 | 0.9132 | 1.1489 | 760 | 800 | - |
| PASEF | 3 | 0.7026 | 0.8879 | 440 | 480 | - |
| PASEF | 4 | 0.9395 | 1.1816 | 800 | 840 | - |
| PASEF | 4 | 0.7289 | 0.9205 | 480 | 520 | - |
| PASEF | 5 | 0.9658 | 1.2142 | 840 | 880 | - |
| PASEF | 5 | 0.7553 | 0.9532 | 520 | 560 | - |
| PASEF | 6 | 0.9921 | 1.2468 | 880 | 920 | - |
| PASEF | 6 | 0.7816 | 0.9858 | 560 | 600 | - |
| PASEF | 7 | 1.0184 | 1.2795 | 920 | 960 | - |
| PASEF | 7 | 0.8079 | 1.0184 | 600 | 640 | - |
| PASEF | 8 | 1.1237 | 1.41 | 1080 | 1120 | - |
| PASEF | 8 | 0.8868 | 1.1163 | 720 | 760 | - |
| PASEF | 9 | 1.0447 | 1.3121 | 960 | 1000 | - |

### Slide 5
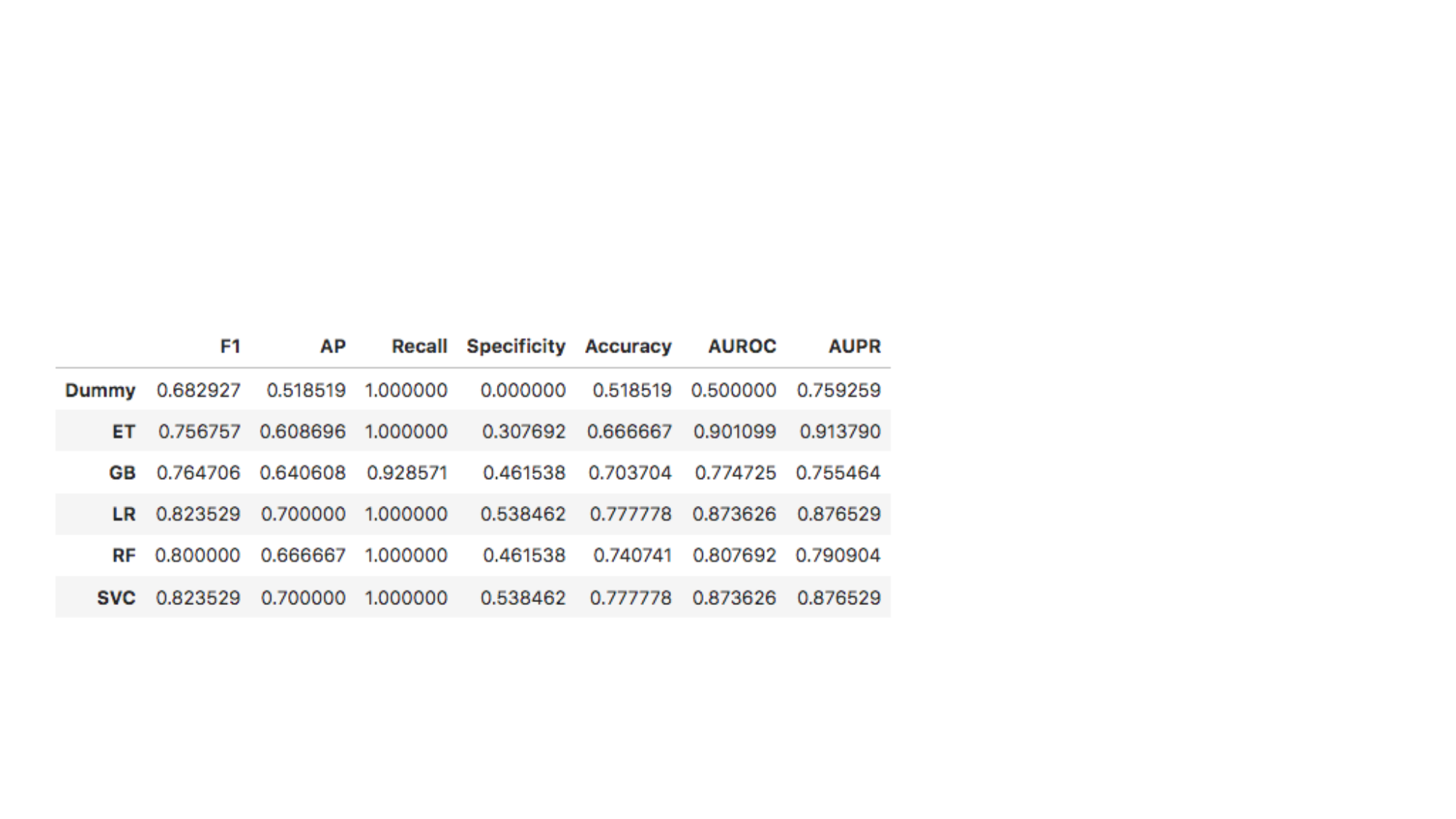
